# Distilling complex evolutionary histories with *shiftPlot*

**DOI:** 10.1101/2022.03.16.484646

**Authors:** Eliot T Miller, Bruce S Martin

## Abstract

Phylogenies form the backbone of many modern comparative methods and are integral components of contemporary science communication. Recent years have seen drastic increases in both the size and complexity of phylogenetic data as computational resources and genetic/trait databases expand. Graphical representations of these massive phylogenetic datasets push against the limits of legibility, often veering closer to artwork than scientific figures optimized to communicate results. While attractive scientific illustrations are certainly a laudable goal, researchers may want to opt for simpler representations to communicate results more concisely. Here, we introduce a new R package, *shiftPlot*, which implements methods for simplifying and plotting phylogenetic comparative data on discrete traits. Specifically, *shiftPlot* automatically finds and collapses clades exhibiting the same character state, effectively creating smaller phylogenies that may be more legibly rendered on standard page sizes. Further, these visualizations more clearly communicate evolutionary dynamics by emphasizing state shifts over tip states. While there are undoubtedly situations where this graphical approach will not be suitable (e.g., continuous traits), we believe *shiftPlot* will prove useful for modern researchers faced with the task of communicating the results of complex phylogenetic analyses.

## Introduction

The derivation of the modern form of the phylogeny, and one of the first graphical depictions of evolutionary relationships, is usually attributed to Charles Darwin (1859). Within a few years, even more visually instructive phylogenies had been derived and used to great effect, as in Ernst Haeckel’s well known tree of plants (Haeckel, 1866; Hoßfeld et al., 2017). This progression towards condensing more information, including trait values and divergence times (Maddison & Maddison, 2009; E. T. Miller et al., 2013; Revell, 2012), across larger trees has continued, supercharged in recent decades by an exponential increase in genomic data and computing power. Today, the pages of many scientific journals regularly contain phylogenies of both epic taxonomic breadth and fine-grained species-level sampling, with trait values exquisitely arrayed across tips of the tree and models of evolution illustrated along branches (A. H. Miller & Stroud, 2022; Williamson & Witt, 2021). Yet, articles are still typically published in standard page formats, often rendering the tip labels and branching patterns of these scientific works of art illegible. While scalable tree viewing software has been used effectively for many years (Letunic & Bork, 2019; Rambaut, 2012; Rosindell & Harmon, 2012), there is currently no feasible way to incorporate such visualizations into scientific papers. Attempts to incorporate uncertainty in our estimates and presentation are indubitably important for both analysis and figure preparation. However, researchers will continue to summarize their results in printed format for many years to come and may wish to consider simpler graphical layouts. It is time to consider when, and how, phylogenetic data visualization can be simplified for improved clarity and communication.

Simpler presentations may be particularly effective in illustrating the distribution of discrete traits across large phylogenies, especially when the trait in question shows a limited number of state shifts, or the shifts themselves are of more interest than tip states *per se*. Tip labels are often far too small and illegible in large phylogenies plotted in standard page format. However, if many tips are collapsed into clades of a single state, effectively creating a smaller phylogeny, trees and trait data may be printed with clade names readable even on a printed page. Furthermore, these simplified phylogenies visually emphasize the timing and pattern of state shifts as opposed to trait data at the tips, more concisely communicating the evolutionary dynamics of interest. For example, Toda et al. (2021) recently used our method, introduced below, to distil 31 shifts in character state across a phylogeny of all bird families (Kimball et al., 2019), allowing for a great deal of clade compression in their figures (Fig. 1). This simplification greatly facilitated communication and discussion of results. While this approach to visualizing discrete trait data on phylogenies may appear rather trivial on the surface, it runs counter to the recent trend of plotting increasingly large phylogenies and trait datasets in formats often impenetrable to understanding.

**Figure 1.**
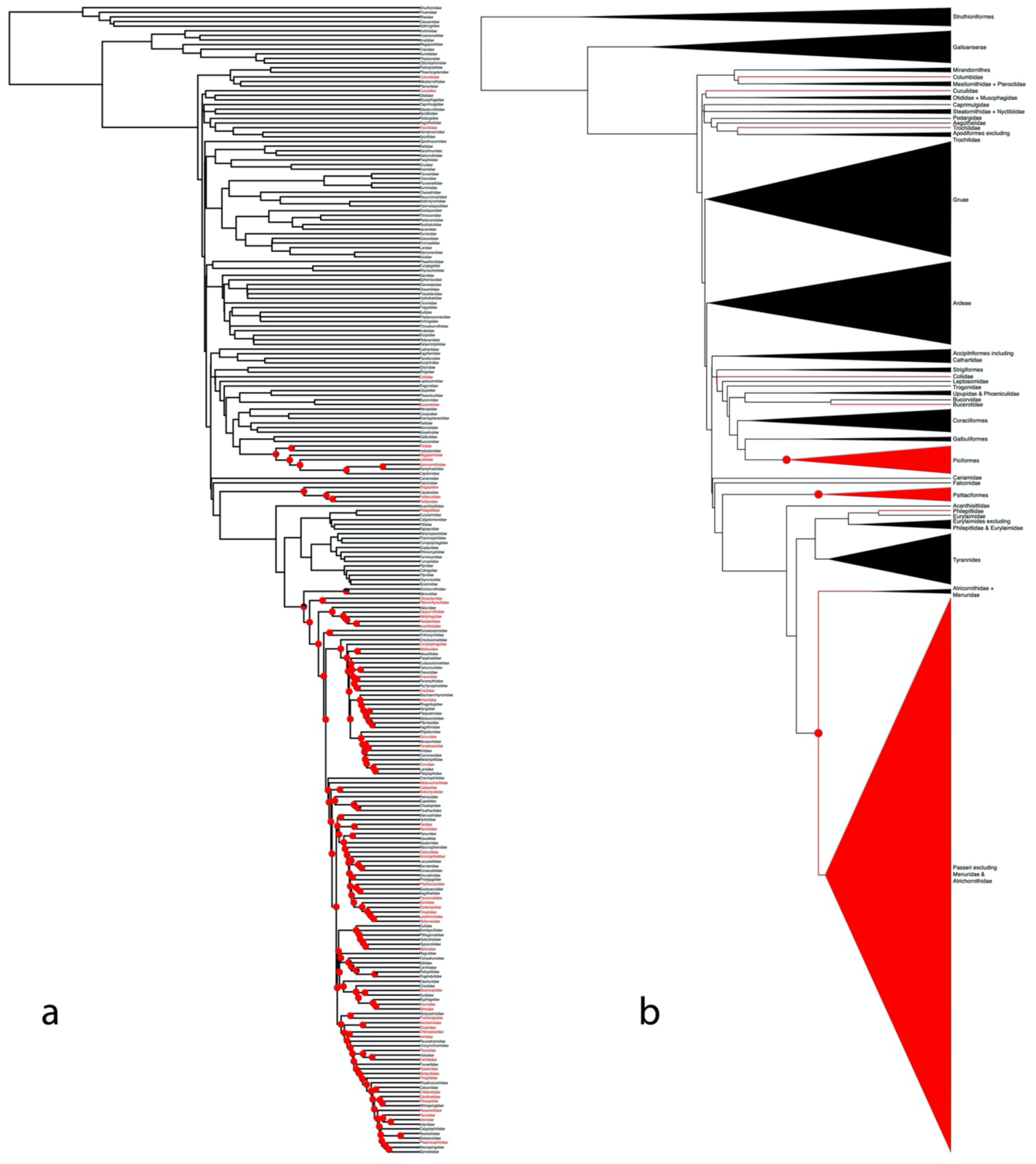
The distribution and inferred origins of sweet taste perception across the avian tree of life. (a) A family-level phylogeny, adapted from Kimball et al. (2019), where internal nodes with greater than 50% probability of possessing sweet taste perception are plotted with pie charts proportional to their inferred state, and tips are coloured according to their observed state. (b) The *shiftPlot* representation of the model. The automatically generated clade names have been replaced with more descriptive and taxonomically accurate names here.

## Implementation

Here we introduce a new R package, *shiftPlot* (https://github.com/eliotmiller/shiftPlot), intended to facilitate visualizations of this nature. The package’s main functionality takes a phylogeny and trait states for all nodes, internal and tip, as input and uses an algorithm to output the maximally condensed phylogenetic representation of the data. The algorithm works by selecting a tip and traversing the tree rootwards, assessing each internal node until it encounters a node where all descendent nodes do not share the same state as the original tip. It then moves back toward the original tip by one node and logs this node as a collapsible entity. All other tips descending from this node are excluded from further consideration. These steps are repeated for remaining tips until all collapsible nodes are identified. This information is fed into an automatic pruning algorithm, which leaves only a single taxon per collapsible clade. The reduced phylogeny can then be expanded to include all original taxa by rebinding removed species as polytomies, setting the stage for using triangles to represent the size of the collapsed clades (Fig. 2).

**Figure 2.**
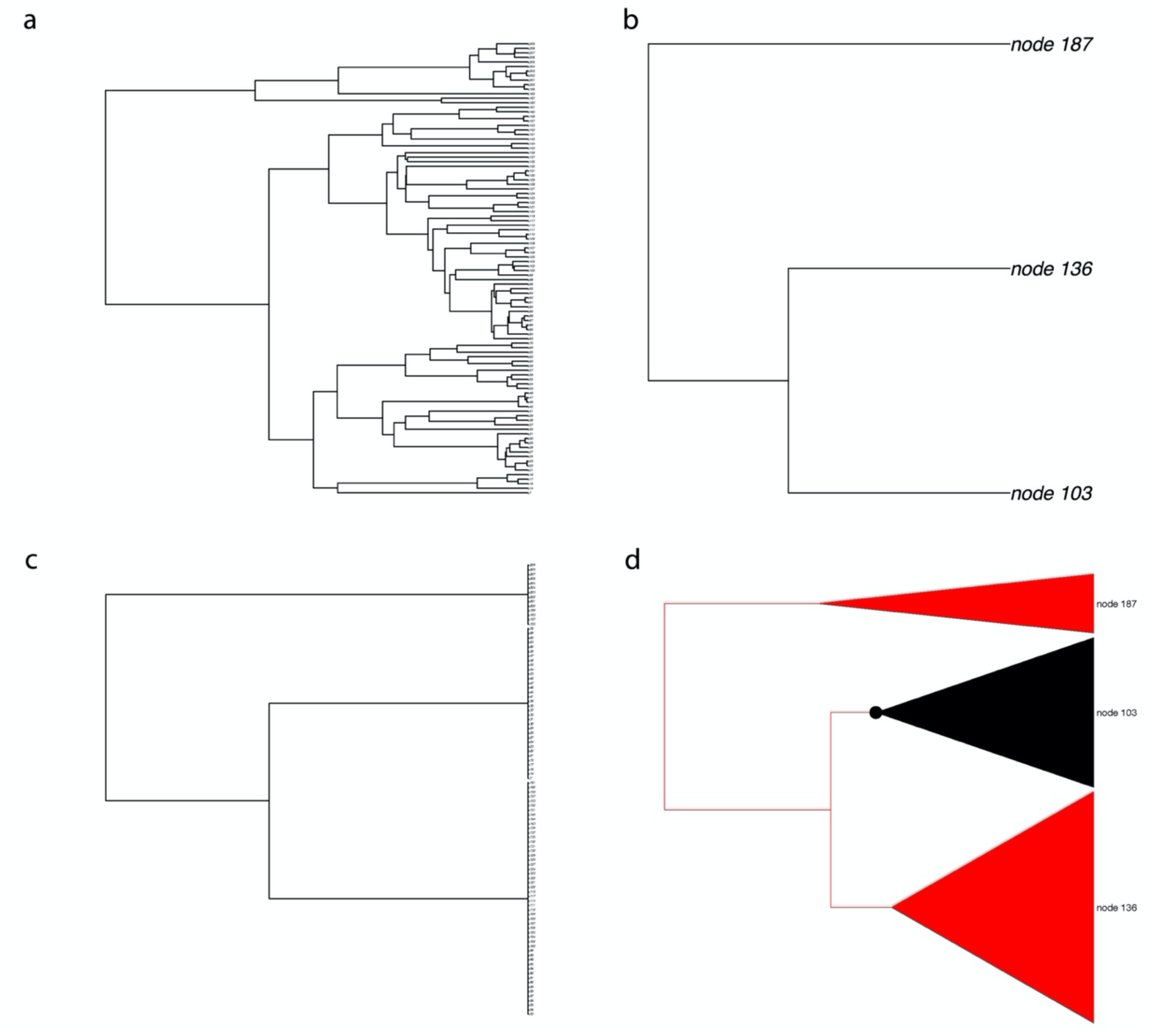
An example to illustrate the way in which *shiftPlot* derives the optimal plot. First, the original phylogeny (a) is optimally collapsed so that it contains the minimal amount of tips which will still permit the evolutionary history to be illustrated (b). Next, the pruned tips are added back in as polytomies (c). Finally, triangles are layered over the top of the polytomy (d). The “point” of these triangles lies at the crown age of the collapsed clade.

The *shiftPlot* package also includes several other functions for facilitating communication of trait evolution, including functions to output details on each change in state (e.g., node and descendent taxa involved in each shift), as well as identifying matching nodes between full and collapsed phylogenies. The latter function works quickly across large trees to identify concordant nodes, and thus is generalizable to phylogenies with differing topologies and compositions, which should prove useful to researchers wishing to identify equivalent nodes across large samples of possibly conflicting trees.

## Discussion

We originally formulated *shiftPlot* as a tool to visualize results directly from *corHMM* (Boyko & Beaulieu, 2021), an R package used for fitting hidden Markov models of discrete character evolution. However, since the package generally requires minimal inputs (a phylogeny and associated node states), it functions across platforms to address the general issue of overly complex phylogenetic data presentations. Indeed, we have recently added functionality to convert stochastic character maps from *phytools* to the relevant *shiftPlot* inputs (Huelsenbeck et al., 2003; Revell, 2012). While the approach to data simplification implemented here will not work for all situations (e.g., continuous trait data), authors may wish to consider opting for simpler, more interpretable phylogeny and trait data presentation using *shiftPlot* when appropriate. We hope this package will inspire others to seek alternative and more general methods for visually condensing phylogenetic and trait data as phylogenetic analyses continue to grow in size and complexity in coming years.

## Acknowledgements

We thank Jacob Berv for insightful comments that improved this manuscript.

## Author’s Contributions

E.T.M. designed, developed, and maintains the software; B.S.M. contributed code. E.T.M. and B.S.M. wrote the manuscript.

## Conflict of interest

The authors have no conflicts of interest to declare that are relevant to the content of this article.

## Data availability

This article is not associated with any original data necessary to archive in conjunction with publication. Example data is included within the R package itself.

## Notes

### Competing Interest Statement

The authors have declared no competing interest.

